# *GoFish*: A low-cost, open-source platform for closed-loop behavioural experiments on fish

**DOI:** 10.1101/2022.04.04.486957

**Authors:** Victor Ajuwon, Bruno F. Cruz, Paulo Carriço, Champalimaud Foundation Scientific Hardware Platform, Alex Kacelnik, Tiago Monteiro

## Abstract

Fish are the most species-rich vertebrate group, displaying vast ecological, anatomical and behavioural diversity, and therefore are of major interest for the study of behaviour and its evolution. Despite this, with respect to other vertebrates, fish are relatively underrepresented in behavioural research. This is partly due to the difficulty of implementing stimuli, manipulanda, and data recording underwater, meaning that this is frequently done with gates to control subjects, physical displays as stimuli, and visual annotation of videos to record data. To overcome these restrictions we developed *GoFish*, a fully-automated platform for behavioural experiments. *GoFish* includes real-time video tracking of subjects, presentation of stimuli in a computer screen, an automatic feeder device, and closed-loop control of task contingencies and data acquisition. The design and software components of the platform are freely available, while the hardware is widely available and relatively inexpensive. The control software, *Bonsai*, is user-friendly and supported by a growing community of users. As an illustration and test of its use, we present the results of 2 experiments on discrimination learning, reversal, and choice in goldfish (*Carassius auratus*). *GoFish* enables the relatively easy implementation of high-throughput tasks and the acquisition of rich behavioural data. Our platform has the potential to become a widely used tool that facilitates complex behavioural experiments in aquatic species.

## Introduction

Behavioural research involves displaying stimuli, detecting subjects’ movements, programming outcomes according to selected contingencies, while recording all of this, including temporal details. This has led to the development and use of conventional experimental platforms that satisfy these needs while ensuring replicability across laboratories. Perhaps the archetype of such a system is the Skinner box designed for pigeons and rodents, that uses manipulanda suitable for those taxa, detecting behaviour by the closing of circuits through key-pecking, lever-pressing, or interruption of light beams. Such systems promote and enhance reproducibility, a critical need in contemporary behavioural research. However, the wide-spread use of a small number of proprietary platforms has, over the years, skewed animal learning and cognition research toward a limited number of study species, including pigeons, rodents, primates, corvids and more recently dogs (Healy, 2019; Shettleworth, 2009) for which such platforms exist. We aim to mitigate this bias by presenting a platform appropriate for fish experiments, that affords all these advantages while being cheap and made of easily sourceable components, as well as being controllable by open source software.

Studying the behaviour of organisms living under water, such as fish, cephalopods, crustaceans, and so on, poses different challenges from those in terrestrial species. In fact, a likely reason why, in spite of fishes’ huge ecological, neuroanatomical and behavioural diversity there is a relative paucity of studies compared to mammals and birds, is the unavailability of standardised programmable experimental platforms that allow for efficient, high-throughput testing and data acquisition with high replicability (Gerlai, 2019; Lieggi et al., 2020). Unlike in most mammal and bird studies, the display of stimuli and delivery of food reinforcers for fish is frequently mechanically executed by an experimenter, increasing temporal variability and vulnerability to observer effects, while restricting scalability. Similarly, data are often recorded by video but annotated visually or digitised at a later time, rather than processing responses in real time so that behaviour can control reward through programmed contingencies. This reflects the fact that most automated operant apparatus that have been developed specifically for fish (Chase & Hill, 1999; Davis & Kenyon, 1971; Horner et al., 1961; Longo & Bitterman, 1959; Talton et al., 1999; Yan & Popper, 1991) were produced decades ago, and hence lack versatility, may require local expertise to operate, and are not sufficiently documented or supported to facilitate easy adoption and wide-spread use.

To address this, we developed *GoFish*, a user-friendly platform for dynamic, fully-automated behavioural experiments on fish or other aquatic organisms. Our platform is inspired by present day behavioural, cognitive and neuroscience experiments that rely on open-source, community based, *DIY-type* solutions for running and developing new experimental paradigms, as well as for processing and analysing the resulting data streams (Akam et al., 2022; Aoki et al., 2015; Bishop et al., 2022; Buscher et al., 2020; Devarakonda et al., 2016; Geissmann et al., 2017; Guilbeault et al., 2021; Gurley, 2019; Kane et al., 2020; Kapanaiah et al., 2021; Lopes et al., 2021; Mathis et al., 2018; Oh et al., 2017; O’Leary et al., 2018; Pineño, 2014; Siegle et al., 2017; Swanson et al., 2021; Walter & Couzin, 2021).

Briefly, our system allows for the display of stimuli on a computer screen placed outside but adjacent to a tank, the detection of the subject’s location in real-time through an overhanging camera, the programming of contingencies between fish movements and the delivery of food rewards, and the automatic recording of data in analysable format. The platform is cheap to build and the design we offer is free and open. Here, we describe the system and present two closed-loop experiments aimed at demonstrating its performance as a research tool. Although we describe an implementation for goldfish, *GoFish* can, in principle, be used with other aquatic species with minimal modifications.

As a proof-of-concept, we show that individual goldfish can be trained to (a) associate a signalled location with food reward and reverse preference appropriately when the contingencies are reversed (*Experiment* 1), and (b) discriminate coloured visual stimuli that switch location between trials (*Experiment 2*).

### The *GoFish* platform

The setup as presently implemented (**Figure 1**) comprises a rectangular prismatic experimental tank (60 x 30 x 36 cm (*length* x *width* x *height*), **Table 1**) with a 17’’ LCD computer screen for stimulus presentation (**Table 1**), placed directly adjacent to the side of the tank where reward pellets are delivered, (**Figure 1a**). Two custom-made, automated pellet dispensers (i.e., feeders (**Figure 1b,c, Table 1**) are clamped onto the upper edge of the tank such that pellets fall on the water surface approximately 2 cm from the closest side of the tank and 2 cm from the screen. Each feeder is placed on either side of an opaque, white acrylic divider (fixed with silicone sealant), running perpendicular to the LCD computer screen, 25 cm into the tank. This partition defines the 2 choice zones of a Y-maze configuration. An overhanging USB camera (1280×720 resolution, **Table 1**) held above the tank records each session (**Figure 1a**). A laptop (**Table 1**) controls task contingencies (stimulus presentation and reward delivery) and video acquisition with a *Bonsai* (Lopes et al., 2015, 2021; Lopes & Monteiro, 2021) custom workflow. A light source (**Table 1**) is placed outside the tank, opposite to the LCD computer screen (**Figure 1a**). The tank is surrounded by opaque styrofoam panels to visually isolate the fish during experiments. The water level is maintained at approximately 15 cm. In the experiments described below, 2 identical experimental tanks were run concurrently, with each fish being tested always in the same tank.

**Figure 1.**
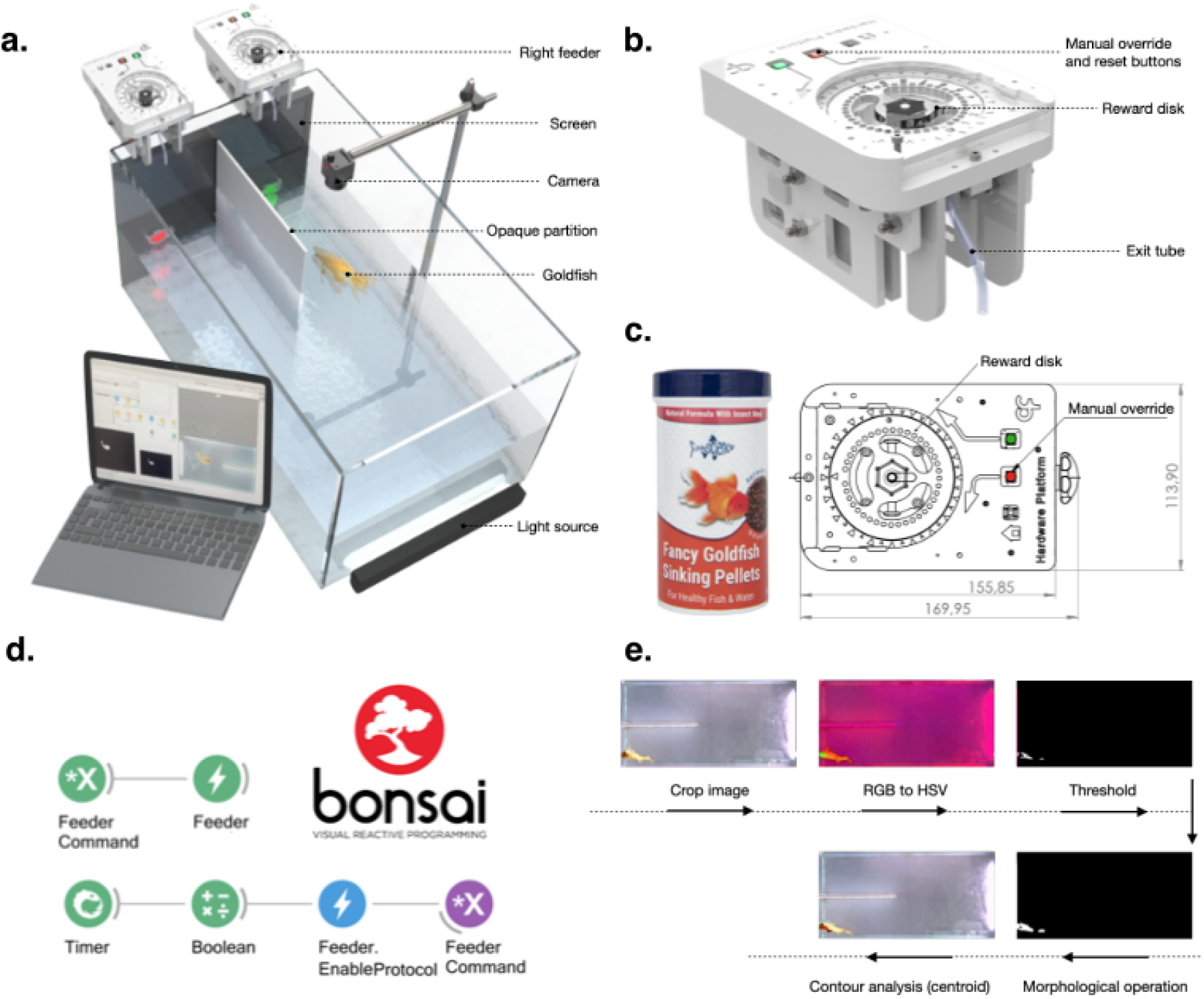
*GoFish* apparatus, feeder control and specifications, and video tracking pipeline. **a.** 3D view of closed-loop operant chamber. Setup includes 2 custom made feeders (see details in c. and e.), computer screen, usb camera and light source. **b.** 3D depiction of the feeder. **c.** Food rewards (left) and detailed top view of feeder reward container disk. **d.** Example Bonsai workflow for custom feeder control. The code implements periodic delivery of food pellets. **e.** Real-time video analysis pipeline, see also Supplementary Videos 1 and 2.

**Table 1.** Parts list. Details of the components used to build one closed-loop behavioural chamber for goldfish learning experiments. Many of the components can be swapped for items of similar functionality to suit particular needs. Feeder parts list and assembly instructions can be found in a dedicated repository (see ***Custom automatic feeder*** section for details). Prices are from early 2021, rounded to the nearest pound.

| **Component Name** | **Supplier** | **Reference code** | **Quantity** | **Unit price** | **Total Price** |
| --- | --- | --- | --- | --- | --- |
| *Clearseal All Glass Aquarium (24x15x12in; LxHxW)* | Goldfish Bowl, Oxford, UK | N/A | 1 | £55 | £55 |
| *WIFIGDS 17 Inch Monitor* | Amazon | B08J7JJD3G | 1 | £81 | £81 |
| *1080P 2MP ELP Web Camera* | ELP | ELP-USBFHD08S-UK | 1 | £72 | £72 |
| *Fish Feeder* | Champalimaud Hardware Platform | FF_GF_SP_V1.1 | 2 | £142 | £284 |
| *IREENUO Aquarium Light (11 inch)* | IREENUO | HDMU-ATL-18A-DE | 1 | £14 | £14 |
| *Laptop computer* | Dell | Latitude 5520 | 1 | £537* | £537 |
| *5mm White Acrylic (300x200x5mm)* | Cut My Plastic | N/A | 1 | £10 | £10 |
| *Betta Clear Silicone 310ml* | Goldfish Bowl, Oxford, UK | N/A | 1 | £13 | £13 |
| *Rankie USB 2.0 Extension Cable* | Amazon | B01KWOBK84 | 3 | £5 | £15 |
| **Combined Total** |  |  |  |  | **£1081** |

\*UK educational price (VAT exempt).

### Pellet dispensers

Design and assembly instructions for laser cut acrylic and 3D printed parts for the pellet dispensers are available from the public repository (https://bitbucket.org/fchampalimaud/device.pump.fishfeeder/).

The instructions include PCB manufacturing plans and specifications, as well as downloadable firmware.

### Stimuli

The potential visual stimuli and their positions are only limited by the monitor employed and its chromatic properties and dimensions. For the experiments described here, the main stimuli were coloured circles (red, green, blue and white, 3.5 cm in diameter, **Figure 2b**) on a grey background, presented with centres positioned 5 cm from the bottom of the tank and 7 cm from each side wall (**Figure 1a**). All stimuli were programmed using custom *Bonsai* (Lopes et al., 2015; Lopes & Monteiro, 2021) and *BonVision* (Lopes et al., 2021) allowing easy generation and manipulation of visual stimuli. Each fish had a randomly assigned unique pair of colour-reward contingencies (**Figure 2b**). We chose colours that have been physiologically (Neumeyer, 1984) and behaviourally (Zerbolio & Royalty, 1983) proven to be discernible by our experimental species (goldfish, *Carassius auratus*). In a pre-experimental, pre-training phase (see details below), we used a white noise rectangle (13.5 x 12 cm, Gaussian: mean=0, variance=10) presented on either the left or right arms of the tank, or in both simultaneously, to signal the imminent delivery of reward in early pre-training, or to signal that reward delivery was contingent on fish swimming to a specific location in later pre-training stages.

**Figure 2.**
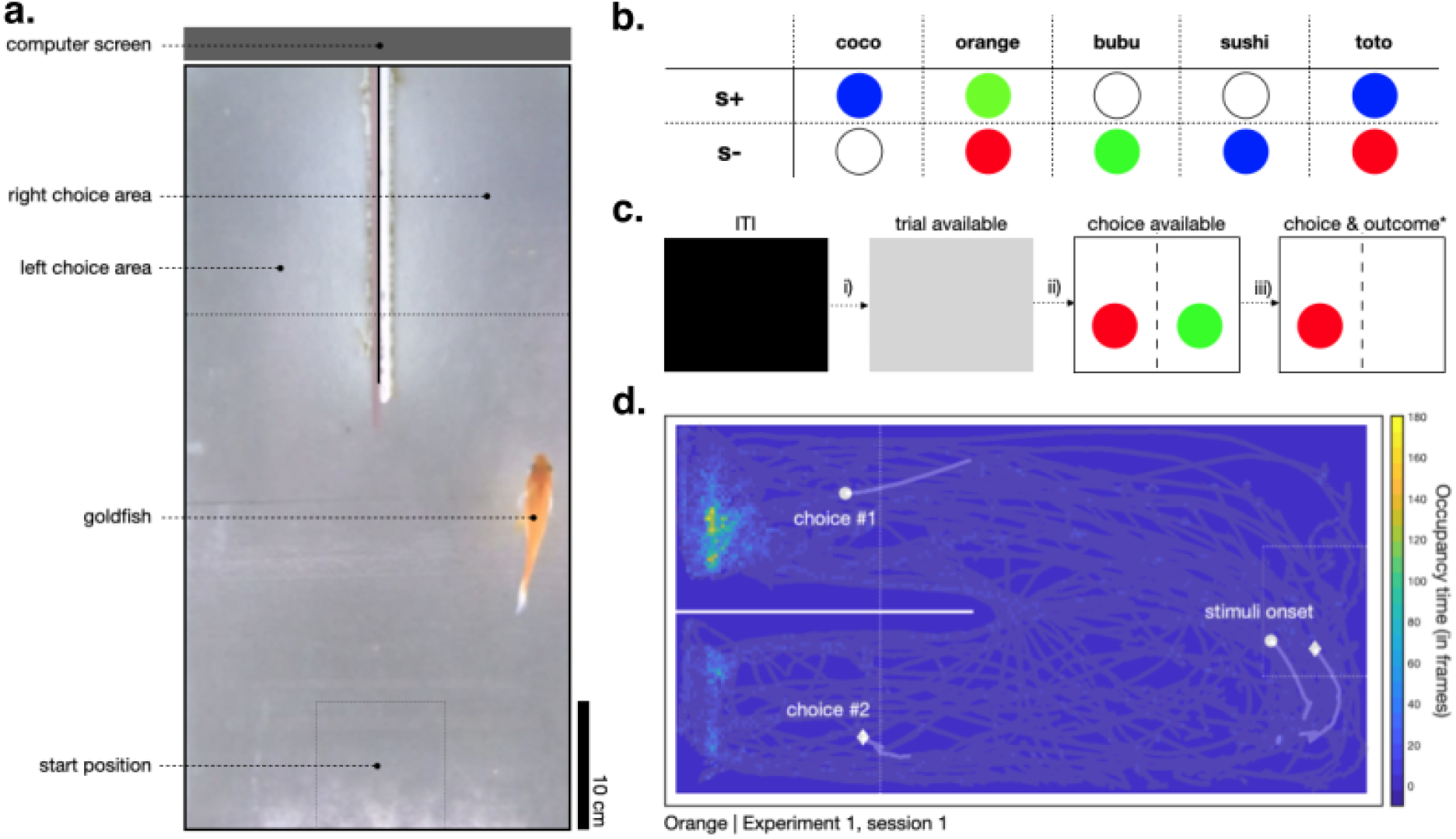
Goldfish were trained to associate colours with food rewards. **a.** Top view of the experimental tank, highlighting start position and left and right choice areas. **b.** Stimuli colour allocation across subjects. **c.** Trial structure: Every trial started with an ITI, signalled by a black screen, drawn from a uniform distribution (min = 20s, max = 40 s), at ITI offset (i), screen becomes grey, signalling fish can move to the start position. As soon as the fish entered the start location, 2 symbols would appear on each side of the screen (ii). Choosing S+ stimulus led to a pellet reward after a 5 second delay, choosing S-started a new ITI. **d.** Grey traces show the position tracks for an example animal and session. Markers depict the position of the fish (i.e., centroid) in the second leading to the stimuli onset and choice, for two example trials (shown with different markers), respectively. The underlying heatmap shows a 2d histogram of occupancy times (in number of frames) for the entire session. Same configuration as presented in a., rotated −90° for presentation purposes.

### Fish tracking

For fish video tracking in real-time we used a colour thresholding method (Monteiro et al., 2021) implemented using custom Bonsai workflow (see example in **Figure 1e**). Video was recorded at approximately 33 fps. Frames were cropped down to include only the inside of the tank and converted to HSV colour space. An HSV threshold was applied to isolate the fish body from the overall white background given by the tank’s bottom. Prior to the onset of the experiment, HSV value ranges were manually set for each fish so as to provide robust tracking in spite of individual differences in fish coloration. The resulting binarized region (pixels are either fish or no-fish) was smoothed and the coordinates of the animals’ centroid were extracted. Occupancy data from a representative experimental session can be found in **Figure 2d**.

### Behavioural task control

Task control was fully automated and implemented using a custom Bonsai workflow (https://github.com/PTMonteiro/GoFish_Ajuwon_etal_2022). Progress through trials was controlled using real-time video analysis of fish movement within the experimental tank. After a variable inter-trial-interval (ITI) fish could advance a trial by swimming into the ‘start zone’. In the main experiment this was a 10 x 10 cm area opposite the rewarded side of the tank that was equidistant to both choice arms (**Figure 2a**). Presence in the start zone after the ITI would trigger stimulus presentation, and a subsequent crossing into either the ‘left choice zone’ or ‘right choice zone’ (15 x 15 cm; **Figure 2a**) would trigger appropriate contingencies. Outside of these epochs (and locations) the fish position had no influence on the unfolding of the task. Note that the ‘start zone’, ‘left choice zone’ and ‘right choice zone’ were not delineated by physical boundary markings, but were defined as specified regions of interest (ROIs) on the video feed corresponding to fixed areas within the experimental tank. Users may wish to make the ROIs visually identifiable to the subjects, as this may influence speed of acquisition. Frames from these ROIs were converted to HSV colour space and a HSV range was applied so as to successfully detect fish. The pixels of the resulting binarized frames from each ROI were summed continuously. Fish entry into the zones was detected from summed ROI pixels exceeding a set threshold (the value of which was adjusted to each subject prior to the onset of the experiment).

### Subjects

Five goldfish ranging in size between 7 - 10cm, (age and sex unknown) participated in the current study. Animals were obtained from a local, commercial supplier (Goldfish Bowl, Oxford, UK).

Fish were housed in groups of two or three, in holding aquaria (60 x 35 x 31 cm; (*length* x *width* x *height*)) where they had access to a rock shelter, pebbles and artificial plants. They participated in experiments 5 times a week on weekdays and fed a total of 24 sinking pellets a day (*Fancy Goldfish Sinking Pellets*, **Figure 1e**). This diet was supplemented with spinach following experiments on the last day of the week and bloodworms the day after. Fish were kept under a 12:12 h light:dark cycle using fluorescent lights. Water was maintained at a minimum of 21°C using an internal heater and independent thermometer (pH: 8.2; ammonia: 0 ppm; nitrite: 0 ppm; nitrate: max. 30 ppm). Partial water changes were conducted at the end of each week and internal filters were cleaned every month. Each holding tank was aerated using an air pump.

For each daily session, fish were transported in a plastic jug to its experimental tank and then back to its holding tank at the end of the session. At the start of each day ~20L of water from all holding tanks were transferred to the experimental tanks in order to keep the environmental conditions as constant as possible. The experimental tanks were cleaned at the end of each week. All animals had experimental experience with unrelated contingencies.

### Pre-training

**Phase 1:** During a 10 min period, fish were allowed to explore and get acclimated to the tanks, previously baited with 12 food pellets throughout. This phase lasted for one day. **Phase 2:** After the ITI (drawn from a uniform distribution: min = 5 s; max = 10 s) during which the screen was black, a white noise rectangle would signal potential food availability in either the left or right choice zones of the tank. Reward was then contingent on fish entering the choice zone signalled with the white noise stimulus. For the first 5 days of this phase there was one session of 12 trials per day and in the following five days, one session of 16 trials per day. Following this, for 3 days fish completed 2 sessions of 12 trials each per day. Rewards were evenly split across both choice zones and allocated randomly. A session ended either when all trials were completed or after 30 minutes. **Phase 3:** After the ITI (drawn from a uniform distribution: min = 20 s; max = 40 s), trial availability was signalled by a grey screen. During this period, fish were required to swim first to the back half of the experimental tank into a ‘start zone’ (i.e., >30 cm, away from the monitor and feeders) to trigger the onset of the white noise stimulus signalling food availability in either the left or right arm. As in the previous phase, reward was then contingent on fish entering a choice zone signalled with the white noise stimulus. This lasted for a minimum of 3 days. Following this, the start zone length was reduced in half (minimum 3 days), and finally to a 10 x 10 cm centred square (minimum 5 days) that was used in the main experiments (**Figure 2a**). There were 2 sessions of 12 trials each per day. To advance through this phase animals had to successfully consume the 12 food pellets within a 1h limit in each daily session. Failure to do so would terminate the training session, with fish returned to their holding tanks. The remaining food pellets would be made available by the end of the day in the holding tanks.

### Experiment 1: Acquisition and reversal of a spatial conditioning

#### Acquisition Phase

Every fish was presented with one daily session of 24 trials. A trial started with an ITI (drawn from a uniform distribution: min = 20 s; max = 40 s) where the screen was black and behaviour had no consequences. The ITI offset was signalled by a grey screen (**Figure 2c**) and from this moment on, entering the start position (**Figure 2a,d**) would trigger the presentation of both visual stimuli (i.e., S+ and S-, see *Stimuli* above) at fixed left/right locations (**Figure 2b,c**, counterbalanced across subjects). Fish made choices by entering one of the two choice zones (**Figure 2a**). Choosing the S+ side resulted in the delivery of a food pellet after a 5 s delay and the onset of an ITI. Conversely choosing the S-side would start a new ITI after a 5 s delay (**Figure 2c**). This experimental phase lasted for 5 days.

#### Reversal Phase

This phase followed the same contingencies as acquisition, except that the rewarded side for each animal (and accompanying stimuli location) was swapped, remaining the same after that. This phase lasted for 7 days.

### Experiment 2: Colour discrimination

In this experiment the rewarded side (and S+/S-stimuli) was randomised on a trial-by-trial basis. To make more correct choices the fish had to follow the S+ and S-signals, rather than acquiring a side preference and reversing it. This experiment lasted for 25 days.

### Data analysis

Real-time video tracking was conducted using a custom *Bonsai* workflow (see *Fish tracking*, above) which also generated a timestamped event list for each session. Preference and movement time (i.e., initiation and response times) data were analysed using custom Matlab code (R2020a, Mathworks) available at https://github.com/PTMonteiro/GoFish_Ajuwon_etal_2022. Statistical analyses were conducted in RStudio (v1.2.5033; The R Project for Statistical Computing, 2018). For statistical analyses, choice proportion data was arcsine square-root transformed to normalise the residuals. One sample, one-sided t-tests against 50% were used to assess performance at group level.

In both experiments repeated measures ANOVAs were conducted to assess the effect of session (to detect learning effects). In *Experiment 2* repeated measures ANOVAs were also conducted to assess the effect of session terciles on trial initiation times and choices (to detect within session satiation or warming up effects). A type-1 error rate of 0.05 was adopted for all statistical comparisons.

### Ethics statement

All experiments were conducted at the John Krebs Field Station and approved by the Department of Zoology Ethical Committee, University of Oxford (Ref. No. APA/1/5/ZOO/NASPA/Ajuwon/Goldfish), and were carried out in accordance with the current laws of the United Kingdom. Animals were cared for in accordance with the University of Oxford’s “gold standard” animal care guidelines. All experimental methods were non-invasive. No food restriction was necessary as fish were fed high palatable pellets during daily-experimental sessions, supplemented by the end of the day in case fish did not eat the minimum daily requirements, and with raw spinach on by the end of the last weekly experimental session. Their diet also included blood worms on weekends. Maintenance and experimental protocols adhered to the Guidelines for the Use of Animals in Research from the Association for the Study of Animal Behaviour/Animal Behavior Society ("Guidelines for the Treatment of Animals in Behavioural Research and Teaching," 2006). On completion, the fish were reintroduced into holding tanks and eventually returned to the supplier.

## Results and Discussion

We illustrate the potential of the *GoFish* closed-loop platform for automated behavioural experiments in two simple learning experiments with goldfish.

In *Experiment 1*, fish (i) controlled the flow of trials by swimming to a start location, which triggered the onset of visual stimuli in two target sites, and (ii) express a choice by swimming to either target ROI, which triggered (or not, depending on choice) a food reward, followed by an intertrial interval, at the end of which the ‘start’ ROI became receptive and a new trial could be started. Multiple-trial sessions took place without intervention of the experimenter. This protocol was used in an acquisition and a reversal phase. The experiment and its results (smooth, significant acquisition and reversal) are similar to those carried out by Kuroda et al. in zebrafish (*Danio rerio;* (Kuroda et al., 2017)). Repeated measures ANOVAs with session as the independent variable confirmed a significant increase in preference for the rewarded side in both the acquisition (F_4,16_ = 3.02, *P* < 0.05), and reversal (F_6,24_ = 13.84, *P* < 0.0001) phases. In the last session of each phase, subjects’ preference for the rewarded side was significantly above 50% (acquisition phase: 88% ± 0.04 (mean ± s.e.m.); one-sample t_4_ = 5.41 *P* < 0.01, reversal phase: 79% ± 0.04; one-sample t_4_ = 5.97 *P* < 0.01)

In *Experiment 2*, reward location was randomised on a *trial-by-trial* basis so that the visual coloured stimuli and the spatial cues were no longer redundant, instead only the former were reliable signals for reward. At group level, fish readily learned to track the location of reward (**Figure 3b**). A repeated measures ANOVA with session as the independent variable confirmed a significant increase in preference for the side displaying the S+ stimulus (F_24,96_ = 2.01, *P* < 0.01). Data from the terminal session show that the average proportion of rewarded choices was 69% ± 0.09. This result was significantly above 50% (one-sample t_4_ = 2.18, *P* < 0.05) even though one of the five fish failed to learn, as shown in **Figure 3**. Since the same subjects were used in both experiments, carry-over effects from *Experiment 1* (where reward site was constant across) may have influenced acquisition of the random alternation protocol in *Experiment 2*.

**Figure 3.**
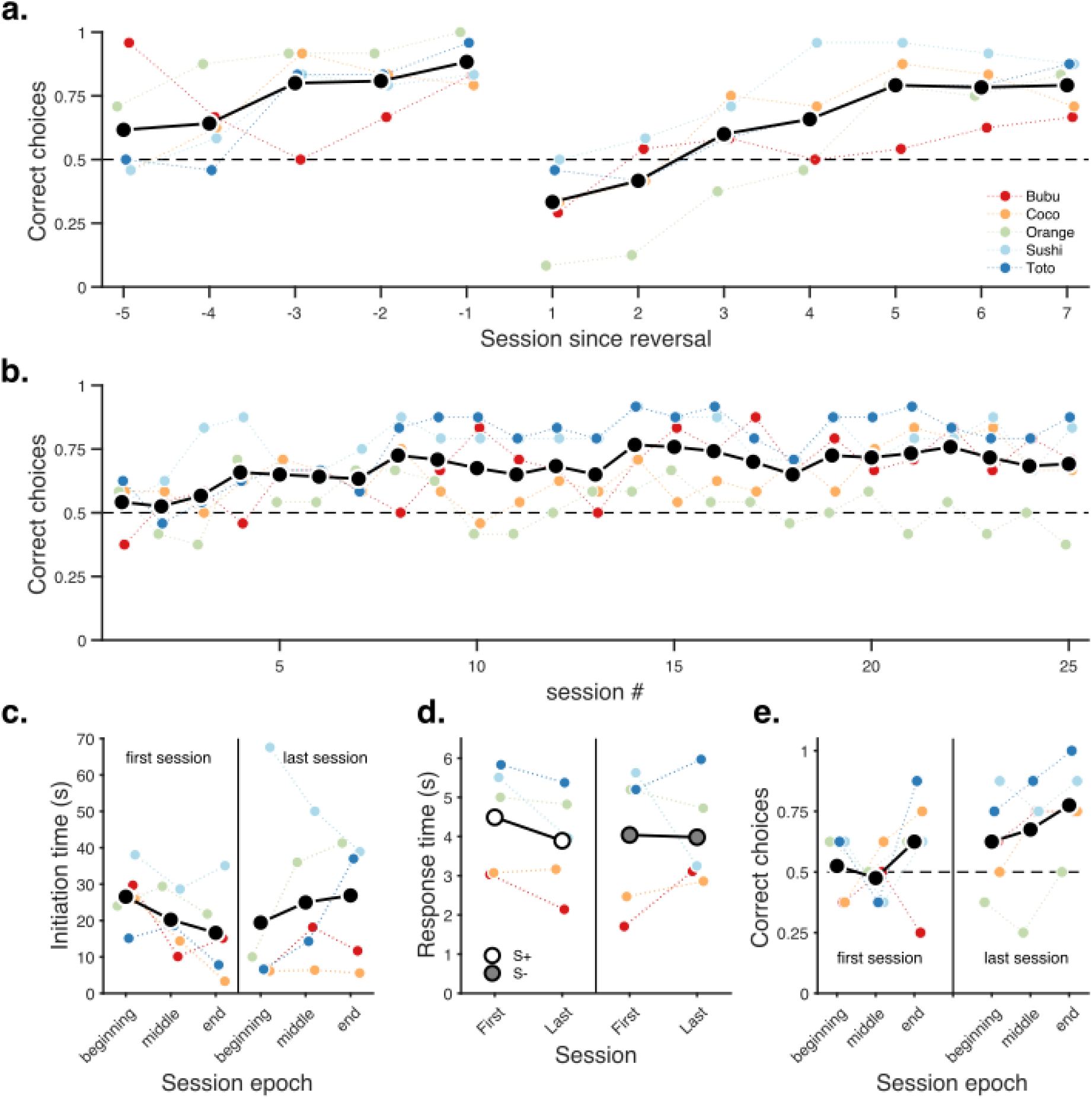
Goldfish learned a colour discrimination task with changing reward/cue/location requirements. **a.** Mean proportion of correct responses for *Experiment 1 (location fixed*) during acquisition and reversal. **b**. Same as a. for *Experiment 2 (trial-by-trial random allocation of stimuli placement*). **c.** Initiation times for the first and last sessions of *Experiment 2*, split into session terciles. **d.** Response times towards S+ (left) and S- (right) stimuli for the first and last sessions of *Experiment 2*, split into the first and last portions of the session, respectively. **e.** Proportion of choices as a function of session epoch (terciles), for the first (left) and last (right) sessions of *Experiment 2*, respectively. In all panels, black (white and grey, for panel d.) markers show group means and coloured markers the mean proportion of correct responses or median initiation or response times, respectively.

In addition to choice data, we used a real-time tracking pipeline for automated detection and recording of fish entry into the regions of interest (start zone, left choice zone, right choice zone). The tracking data gives direct access to relevant behavioural metrics, such as trial initiation time (i.e., the time animals took to be detected in the start zone following ITI offset; **Figure 2c** - ii) and choice response times (i.e., the time from starting a trial to entering one of the choice zones; **Figure 2c** - iii).

As a metric for learning and motivational changes, we compared initiation times between the first and last sessions of *Experiment 2* (**Figure 3c**), but found no significant differences (paired t_4_ = −0.86, *P* = 0.44)

In addition to choice proportion, we measured choice response times. This variable can be extremely informative: in previous studies and protocols it has been found that response times on both single-option and choice trials can be at least as informative regarding preferences and choice mechanisms as choice proportions (e.g., Monteiro et al., 2020). Overall, we found no significant differences in response time between trials in which fish chose correctly or incorrectly (**Figure 3d,** first session: paired t_4_ = 1.61, *P* = 0.18, last session: paired t_4_ = 0.29, *P* = 0.44)

Finally, we explored whether the proportion of correct choices varied within sessions by checking for trends across terciles of sessions. Such effects can occur if there are ‘warming up’ or satiation effects. Once again, we found no significant effects either early or late in training as revealed by repeated measures ANOVAs with session tercile as the independent variable (**Figure 3e,** first session: F_2,9_ = 2.29, *P* = 0.16, last session: F_2,9_ = 0.096, *P* = 0.91).

In summary, as a *proof-of-concept* demonstration for *GoFish*, a fully automated, closed-loop, and open source experimental platform, we show that goldfish can reliably learn to i) associate a fixed location with reward ii) reverse their preference when the rewarded location changes, and iii) associate colours with reward contingencies. We also present temporal data because, although no significant effects were found in this sample study, they illustrate what can be measured and suggest strategies for analysis.

## General Discussion

*GoFish* is a new, user-friendly platform for dynamic, fully-automated behavioural experiments that facilitates high-throughput, highly reproducible behavioural research in fish or other aquatic organisms. *GoFish* is open-source, inexpensive, simple to assemble, highly adaptable, and should be supported by a growing community of users. As a generic experimental platform, *GoFish* provides advances and improvements over more common experimenter-controlled setups currently used in research on fish behaviour and cognition. These improvements fall into four major categories: 1) methodological, 2) welfare, 3) scientific, and 4) educational.

Methodologically, the platform reduces the potential for unintended bias in experimenter-run tests, which are harder to run blindly. Also, due to the fact that tasks become fully automated, it reduces the chance of human errors. Moreover, in combination with the automation, its low cost (**Figure 1a**, **Table 1**) opens the possibility of testing multiple animals in parallel. Such standardisation across setups and subjects increases efficiency and contributes to reducing inter-individual variability, ultimately contributing to a general refinement of procedures.

Methodological refinements will likely result in the reduction of the numbers of experimental animals used. Moreover, eliminating the experimenters’ presence during data collection reduces noises, shadows or other uncontrolled environmental changes thereby reducing subjects’ stress levels, and improving subjects’ welfare.

*GoFish* improves reproducibility, due to standardisation, and highlights the importance of *low-cost open-source* tools for the advancement of science as a whole - the fact that all components (including software) are *open-source* should afford community-based long term further development. Enabling easier automated extraction of a wider range of behavioural metrics should enrich the description of behaviour.

We stress that the experimental configuration presented here (a Y-maze setup for two alternative forced choice reversal learning and colour discrimination tasks) is a *proof-of-concept* but that *GoFish*’s basics could be used without configural changes in multiple other experimental paradigms e.g., quantity discrimination experiments (Potrich et al., 2022; Schluessel et al., 2022), behavioural timing (Talton et al., 1999), foraging (Aw et al., 2009; Newport et al., 2021), object recognition (Newport et al., 2016), and navigation (Burt de Perera & Holbrook, 2012; Newport et al., 2016). *GoFish* could also be used to implement experiments using a range of set-ups differing to that reported here (e.g., open field and maze configurations that could employ a greater number of screens and/or feeders than we have). It is worth noticing that *GoFish* could be used to present stimuli, in other sensory modalities: instead of using computer screens for visual stimuli presentation, *Bonsai* affords a large pool of interaction possibilities (e.g., adding a range of sensors and/or actuators, and sound libraries for auditory stimulus generation (Lopes et al., 2021). Moreover, within *Bonsai*’s framework, our tracking routine, based on colour thresholding, could be extended to implement marker-less (Kane et al., 2020) and multi-animal tracking (Guilbeault et al., 2021). Furthermore, our automatic feeder could easily be modified to use other regular-shape rewards by 3D printing a different reward disk (**Figure 1c**) (Arce & Stevens, 2022; see also Oh et al., 2017). With the present dimensions, the maximum number of rewards between re-fills is 40, which may be limiting for some applications. However, this number depends on the size of individual rewards, which may vary depending on the particular application of the feeder.

Finally, we note that the low price and scalability of the system makes it suitable for hands-on practical experiments and projects in education contexts (e.g., undergraduate projects, summer courses). It could be used for teaching basic animal learning, experimental methods for behavioural research, and data processing (i.e., video tracking) and visualisation.

*GoFish* is a fully integrated, adaptable platform designed to facilitate the easy implementation of complex behavioural protocols in aquatic species. We hope that our platform accelerates the pace of behavioural research in a range of species that otherwise have been relatively underutilised in comparative and cognitive research programmes.

## Acknowledgements

We thank Adelaide Sibeaux for technical advice and husbandry training, Theresa Burt de Perera, Cait Newport, Christine Soper and all the Wytham Field Station staff for logistical support.

## Author Contributions

VA, AK, and TM conceptualised and designed the experiments; VA and TM built the set up; VA, BC and TM wrote the Bonsai workflow; VA and TM collected the data; VA, BC, AK and TM analysed the data; PC and CFSHP designed and built the feeders; VA and TM wrote the first draft of the paper; all authors edited and reviewed the manuscript.

## Data and Code accessibility

Data is available as a supplementary file (**Data file 1**) and code for running the task and data analysis can be accessed at https://github.com/PTMonteiro/GoFish_Ajuwon_etal_2022.

## Supplementary materials

**Data file 1.** *“Figure3.xlsx”* contains all data presented in **Figure 3**. Each tab corresponds to a panel in the figure. *“panelA”* and *“panelB”* tabs include mean proportion data by session (rows) and animals (columns) for *Experiments 1* and *2*, respectively; *“panelC”* tab contains individual median initiation times (rows) for the first and last sessions’ terciles of *Experiment 2* (columns A:C and D:F, respectively); *“panelD”* tab contains individual median response times (rows) toward stimulus S+ during the first (column A) and last session (column B) of *Experiment 2*, and the corresponding data for responding towards stimulus S-in columns C:D, respectively; *“panelE”* tab contains individual choice data (columns) for the first and last sessions of *Experiment 2* concatenated vertically (24+24 trials), with ones indicating an S+ choice, and zeros otherwise. To generate Figure 3, run *“Figure3_forShare.m”* available at: https://github.com/PTMonteiro/GoFish_Ajuwon_etal_2022.

**Video 1.** Example real-time video analysis pipeline for a representative animal and session.

**Video 2.** Example of *Bonsai* tracking for a representative animal and session.

